# A novel method for detection of Cas9 gRNAs using a fluorophore-labeled DNA oligo

**DOI:** 10.1101/2025.02.21.639318

**Authors:** Ranmal Avinash Bandara, Zhichang Peter Zhou, Ziyan Rachel Chen, Rongqi Duan, Alan Richard Davidson, Amy P Wong, Jim Hu

**Author notes:** Correspondence: Jim Hu.

## Abstract

Clustered regularly interspaced short palindromic repeats (CRISPR) associated protein 9 (Cas9) has become one of the most important gene editing tools. CRISPR guide RNA (gRNA) serves as an important component in guiding Cas proteins to a target site in gene editing processes. For effective gene editing or site specific gene integration, levels of gRNA present in a target cell are critical. Especially, for prime editing, even the ratio of different gRNAs present in target cells has a major effect on gene editing efficiency. Therefore, having a convenient, highly sensitive method for direct detection of gRNAs can improve editing design and optimize targeting efficiency. Here, we have developed a convenient, highly sensitive method for direct detection of gRNAs, which may be used to improve editing design and optimize targeting efficiency.

## Main text

CRISPR/Cas9 is a versatile genome editing tool that has the potential to be used to cure many genetic diseases. The system works via a gRNA interacting with the Cas9 protein to form a complex that binds to a specific DNA sequence^1^. The site-specific DNA binding feature of the Cas9 system can be utilized in a variety of ways to correct gene mutations or to regulate gene expression. First, the Cas9 protein can make a site-specific double stranded break (DSB) that is mainly repaired by homology-directed repair (HDR) or non-homologous end joining (NHEJ). Both of these pathways can be used for gene editing or gene insertion when proper donor DNA is present^1^. Second, the Cas protein can be modified into a nickase, which creates a site-specific nick. When fused with other proteins, such as adenosine deaminase, cytosine deaminase, or reverse transcriptase, the modified Cas9 protein can be used for base editing^2^ or prime editing^3^. While cytosine base editors (CBEs) and adenine base editors (ABEs) allow single-nucleotide changes at target sites specified by the gRNA, prime editors, with the gRNA modified to include a template sequence for reverse transcription, can offer single nucleotide substitution as well as short deletions or insertions^3^. The recently improved prime editing systems include three gRNAs: an engineered pegRNA (epegRNA) to locate the target site for initiation of the prime editing, a nicking gRNA to nick the unedited DNA strand to enhance the editing efficiency, and a dead single gRNA (dsgRNA) to recruit the Cas9 protein near the editing site to enhance editing efficiency^4^ . In addition, the Cas9 protein can be fused with a transposase, such as in the Find and Cut-and-Transfer (FiCAT) system, to perform site specific gene integration^5^. Finally, a modified Cas9 protein can serve as a site-specific DNA binding protein to regulate gene transcription by fusing with a transcription activator or inhibitory motif^6^. In all the above gene editing or gene regulation approaches using Cas9, proper levels of gRNA presence are essential for targeting efficiency. In the case of prime editing, even the ratio of gRNA expression is critical; the epegRNA level should be at least 3-fold higher than the nicking gRNA or dsgRNA^4^. Currently there is no reliable, convenient method to precisely measure levels of gRNA expression. In this study, we developed a rapid, sensitive, cost-effective and non-radioactive method for gRNA detection.

To design such a method, we considered using a 5’-Cy5.5-labelled DNA oligo to probe gRNAs through in-solution hybridization and detecting the hybridization products on a native polyacrylamide gel with a fluorescence-detection-capable imager, the Licor Odyssey. For the DNA oligo design, we decided to select a sequence that targets a common region of a Cas9 gRNA so that this universal probe can be used to detect gRNAs for different Cas9 target sites and for Cas9 gRNAs used for different applications, including Cas9-mediated gene integration, base editing, prime editing, and Cas9-mediated gene regulation (Fig. 1A).

**Figure 1.**
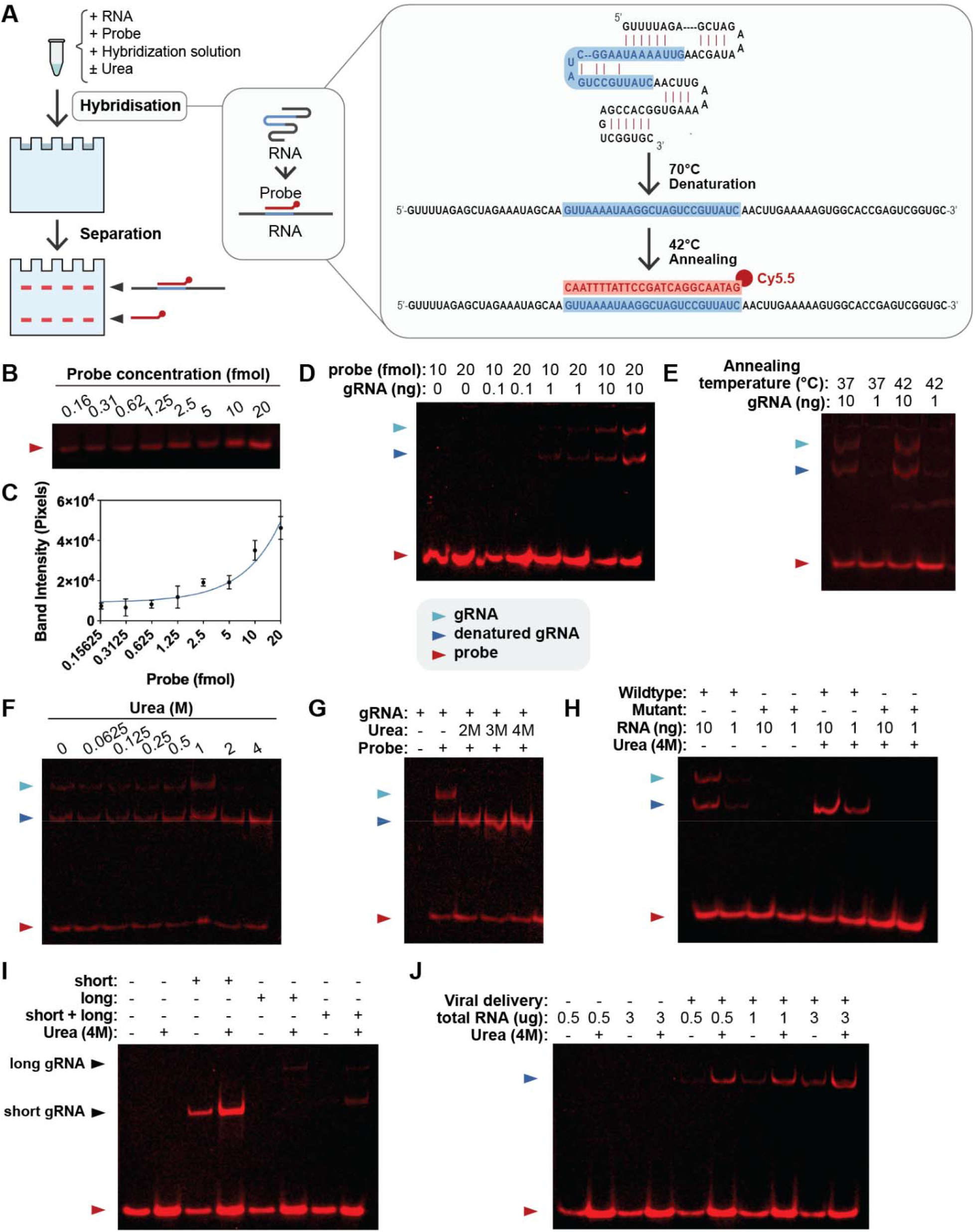
Method design, probe sensitivity test, Suppressing RNA secondary structure and enhancing the detection sensitivity with urea. (A) Diagram showing experimental procedure overview. The red and blue sequences represent the probe and the probe binding region on gRNA, respectively. The red dot represents the Cy5.5 fluorophore. (B) Cy5.5 probe detection using Licor imager after native gel electrophoresis. Different amounts of indicated Cy5.5 probe were loaded onto a native polyacrylamide gel using the Blue Juice dye and run on native polyacrylamide gel for 40 minutes. (C) ImageJ analysis was done on gels (N = 3) as seen on Fig.1B. (D) gRNA detection using Cy5.5 probe. *In vitro* transcribed gRNA was hybridized in solution with 20 fmol of Cy5.5 probe. The hybridization products were separated on 10% native polyacrylamide gel and imaged using the Licor imager. (E) Detection of gRNA in different annealing temperatures. (F, G) Effect of Urea on gRNA detection. 10 ng of gRNA was incubated with 20 fmol of probe as described. Urea was added to indicated concentration and incubated.(H) Absence of detection by mutating the probe binding sequence in the gRNA. (I) Detection of gRNA in 5 µg of total RNA from mammalian cells. Short, long and short plus long indicate total RNA from HEK293T cells transfected with plasmids expressing short, long, or short plus long gRNA, respectively. (J) Detection of gRNA in mammalian cells with viral delivery. 50 MOI of CRISPR/Cas9 viral vector was added to 116 cells.

We first tested the probe sensitivity (see supplementary Fig. 1). To ascertain the probe sensitivity in a polyacrylamide gel is similar to what we observed, different concentrations of the probe were run on a 10% native polyacrylamide gel. As shown in Fig. 1B and 1C, the probe was detectable from 0.16 to 20 fmol. Following confirmation of the high sensitivity, we determined whether our probe can be used to detect gRNAs. We first chose *in vitro* transcribed gRNA for the initial test because its high purity allows for greater specificity and accuracy as compared to gRNAs expressed from cells. To determine the minimal amount of guide RNA (gRNA) that could be detected, we used 10, 1, and 0.1 ng of *in vitro* transcribed gRNA for solution hybridization (see online methods and materials). We observed that 10 ng of gRNA generated the highest signal whereas 1 ng of gRNA showed much lower level of signal (Fig. 1D).

To optimize the annealing temperature for hybridizing the probe and gRNA, the mixture of probe and gRNA was heated to 70 ^o^C for 3 minutes and then cooled down to either 37 ^o^C or 42 ^o^C for 15 minutes. As shown in Fig. 1E, the 42 ^o^C annealing temperature yielded a stronger signal compared to 37 ^o^C. From here, we used the 42 ^o^C annealing temperature for the remainder of our experiments. In this test, we also observed two bands of the hybridization products due to the secondary structure of the gRNA (Fig. 1E). We envisioned that if we could eliminate the secondary structure and allow the hybridization product to run as a single band, we would gain sensitivity in detecting gRNAs in total RNA samples from mammalian cells, especially when multiple gRNAs with different sizes are present.

We used urea to disrupt the secondary structure since urea is known to disrupt hydrogen bonds and interact with nucleotides. We found that 2 M urea started to have a major effect on the gRNA secondary structure. Inclusion of 4 M urea disrupted completely the RNA secondary structure resulting in single band of hybridization product with stronger fluorescent signal without separation of the probe from gRNAs (Fig. 1F and 1G). To ascertain the probe specificity, we designed a guide RNA with the probe binding site mutated and we showed this RNA was not detected by the probe (Fig. 1H).

To determine whether gRNAs from total RNA isolated from mammalian cells could be detected, we transfected HEK293T cells with plasmids expressing gRNAs from the U6 promoter. We found gRNAs can be detected in 5 µg of total RNA from cells expressing single or double gRNAs (Fig. 1I) despite different levels of expression (explained in Supplementary information). We also analyzed the magnitude of urea effects on gRNA detection with the result discussed in Supplementary information as well. Finally, we found that gRNAs from 116 cells transduced with Helper-Dependent Adenoviral (HD-Ad) vector^7^ expressing Cas9 and gRNA can be detected with less than 1 µg of total cellular RNA, indicating the high sensitivity of the method (Fig. 1J).

In summary, these experiments demonstrated the successful development and the utility of a novel technique for gRNA detection. This method is expected to have a wide range of applications for assessing levels of gRNA expression, such as Cas9-mediated gene integration and gene editing as well as gene regulation. Due to differences in target sites and/or in reverse transcription template sequences used in prime editing, different gRNAs may have different levels of expression. This may lead to differences in gene editing efficiency. Our method of gRNA detection can be used for assessing gRNA expression in all these cases and therefore may facilitate the development of genetic approaches for gene therapy applications.

## Author contributions

J.H. conceived this research. R.A.B. performed all the transfections and transductions, RNA isolation, in-solution hybridizations, running of the gels, imaging of the gels, and analysis. Z.P.Z. invitro transcribed the RNA used. Z.R.C. created figure 1A and labelled all figures. R.D. Cultured HEK293T and 116 cells. A.R.D. and A.P.W contributed a critical review, suggested experiments and polished the manuscript. J.H., R.A.B., Z.P.Z., and Z.R.C. wrote the manuscript.

## Data availability

Data is available upon request.

