## Supplementary information for "A novel method for detection of Cas9 gRNAs using a fluorophore-labeled DNA oligo"

**ONLINE MATERIALS**

**Methods and Materials**

Oligo probe

The 5’-Cy5.5 conjugated DNA oligo used in this study was ordered from IDT (Coralville, Iowa). In this probe, a single Cy5.5 fluorophore was attached to the DNA oligo (GATAACGGACTAGCCTTATTTTAAC).

*In vitro* generation of guide RNAs

To make gRNA *in vitro*, we constructed DNA templates containing a cassette, (T7 promoter)- (gRNA target site)-(gRNA scaffold), using PCR. We used the common Cas9 guide RNA scaffold (5’- GTTTTAGAGCTAGAAATAGCAAGTTAAAATAAGGCTAGTCCGTTATCAACTTGAAAA

AGTGGCACCGAGTCGGTGC-3’) and a target site, which locates in the GFP coding sequence (5’-GCC GTC CAG CTC GAC CAG GA-3’). Purified PCR product was used as a template for *in vitro* transcription of gRNA. An *in vitro* transcription kit (NEB, E2040S) was used following the manufacturer’s instructions. Specifically, the reaction was assembled in 20 µL with the following composition: purified PCR product (1 µg), ATP (2 µL, 100 mM), GTP (2 µL, 100

mM), CTP (2 µL, 100 mM), UTP (2 µL, 100 mM), DTT (1 µL, 0.1 M), RNase inhibitor (1 µL;

NEB, M0314S), nuclease-free water, reaction buffer (2 µL), and T7 RNA polymerase mix (2 µL). The reaction was incubated at 37 °C for 16 hours. Subsequently, 28 µL of nuclease-free

water and 2 µL of DNase I were added to the reaction and incubated at 37 °C for 15 minutes. The gRNA was purified using an RNA cleanup kit (NEB, T2050L) following the manufacturer’s instructions.

To generate a negative control gRNA with the targeted region mutated, we ordered primers with the mutated scaffold which prevents binding of the scaffold by the probe. The probe binding region was replaced with the sequence 5’- CTTATTAAGACTCCAACCGAGAGG – 3’.

Cell transfection and transduction

For total RNA isolation from cells with gRNA expression from plasmids, HEK293T cells cultured in DMEM (10% FBS, 1% Penicillin-Streptomycin, 1% glutamine) at 70% confluency were transfected with 6 µg of plasmid DNA using jetPRIME transfection reagent. Cells were cultured for 2 days before RNA isolation. Three plasmids were used in this study and the guide RNAs expressed from the U6 promoter in these plasmids share the common scaffold sequence as the *in vitro* transcribed gRNA mentioned above. The size of the short gRNA only plasmid is 12474 bp and the gRNA expressed from the plasmid is 101 bases in length. This plasmid also expresses a Cas9 protein. The size of the long gRNA only plasmid is 2354 bp and the gRNA expressed from this plasmid is 176 bases in length. The double gRNA plasmid expressing both short and long gRNAs is 12884 bp, which also expresses a Cas9 protein.

For total RNA isolation from cells with gRNA expression from a viral vector, 116 cells cultured in MEM (10% FBS 1% Penicillin-Streptomycin) were transduced with a helper-dependent adenoviral vector expressing Cas9 and a gRNA (HD-Ad-LacZ) targeting the GGTA1 locus^1^.

This vector also carries a donor gene expression cassette expressing the LacZ gene, and it was designed to target the swine GGTA1 locus which encodes a-1, 3-galactosyltransferase that is expressed in many mammals. However, in humans, this gene is a transcribed pseudogene. We use this viral vector to demonstrate the levels of gRNA expressed in mammalian cells. Cells were seeded into a 6 cm plate cultured until 70%-80% confluency before transduction with HD- Ad-LacZ at 50 MOI (multiplicity of infection) as described^1^. Total RNA was isolated two days post transduction.

RNA isolation using TRIzol

Total RNA isolation was performed with TRIzol (ThermoFisher, Mississauga, Ontario, Canada, Cat#: 15596026) according to manufacturer’s guideline. Briefly, 0.4 ml of TRIzol was added to a 6 cm dish and the lysate was transferred to an Eppendorf tube. Following incubation at room temperature (RT) for 5 minutes, 0.1 ml of chloroform was added. The tube was then vigorously shaken for 15 seconds and incubated at RT for 2 minutes. The samples were centrifuged at 12,000 g for 15 minutes at 4 ^o^C. The top layer was transferred to a fresh tube and 0.25 ml of isopropyl alcohol was added. The samples were incubated at RT for 10 minutes and centrifuged at 12,000 g for 10 minutes at 4 ^o^C. The supernatant was removed and 500 µl of 75% ethanol was added. The samples were vortexed and centrifuged at 7,500 g or 9,500 rpm for 5 minutes at 4 ^o^C. The pellet was dried and dissolved in 50 µl of RNase free water.

In-solution hybridisation

RNA was combined with 1 µl of 10X hybridisation solution (1500 mM NaCl, 500 mM Tris-Cl at pH 7.5, 10 mM EDTA), Cy5.5 probe (20 fmol), with or without urea (4M). The total volume was brought to 10 µl with water. Samples of RNA and probe mixture were heated at 70 ^o^C for 3 minutes followed by incubation at 37 ^o^C or 42 ^o^C for 15 minutes and then the temperature was brought down to 4 ^o^C.

Native polyacrylamide gel

To make a 10% non-denaturing polyacrylamide gel, a total volume of 18 ml of polyacrylamide solution (for two gels) is prepared with 5.8 ml of 30% Acrylamide/Bis solution 29:1 (Bio-Rad, Mississauga, Ontario, Canada, Cat#: 1610156), 1.8 ml of 5X TBE buffer, 10.2 ml of water. The polymerization was started by adding 140 µl of 10% ammonium persulphate (Bio Basic, Markham, Ontario, Canada, Cat#: AB0072) and 12 µl of TEMED (BioShop, Burlington, Ontario, Canada, Cat#: TEM001.25). The samples mixed with Blue Juice (Invitrogen, Vilnius, Lithuania, Cat#: 10816015) were run in the dark on the native polyacrylamide gel for 40 min or 1 hour at 100 V.

Imaging equipment

The Licor Odyssey FC at the Imaging facility at the Hospital for Sick Children was used. The 700 channel (685b nm laser) was used for all imaging.

### SUPPLEMENTARY INFORMATION

1) The need for novel guide RNA detection methods

Many approaches of gene editing or gene regulation are based on the CRISPR/Cas9 system. In all these approaches, guide RNAs (gRNA) are required to bind to the Cas9 protein for positioning the RNA/protein complex to a specific chromosomal site.

Although the gRNA scaffold can be the same, different gRNAs are different in their targeting sequence and 3’ modifications^1^. Guide RNAs driven by different promoter may express different levels. Currently there is no convenient method for assessing levels of gRNA expression. Traditional northern blotting requires radioactive isotope and is time- consuming. Because the size of gRNAs is small, the random hexamer primers may not allow different regions of the entire gRNA to be equally reverse-transcribed, leading to inaccuracy in measurement with real-time PCR.

2) Rationale for using Cy5.5 as a fluorophore

We decided to use a Cy5.5-labeled oligo because of the following considerations: First, Cy5.5 is a highly sensitive, near-infrared fluorophore and has low background autofluorescence from biological samples. This is the key requirement for the method development. Second, Cy5.5 is more photostable compared to shorter-wavelength fluorophores, such as Cy3 and Cy5. This allows longer exposure time during the scan to increase the sensitivity of detection. Third, the high affinity of the RNA-DNA base- pairing will allow a short time for hybridization, and the hybridized products will remain stable during gel separation. Finally, using the single fluorophore-labeled oligo for gRNA

detection is a direct approach that can accurately reflect the actual levels of gRNA without the need for amplification such as in real-time RT-PCR.

3) Testing probe sensitivity

To investigate the probe sensitivity, the Cy5.5-labeled oligo was diluted to different concentrations at femtomole (fmol) levels, and 30 µl from each concentration were spotted onto a nitrocellulose membrane and imaged with Licor Odyssey FC in our Imaging facility. We found that the probe was detectable from 0.1 to 10 fmol (supplementary Fig. 1).

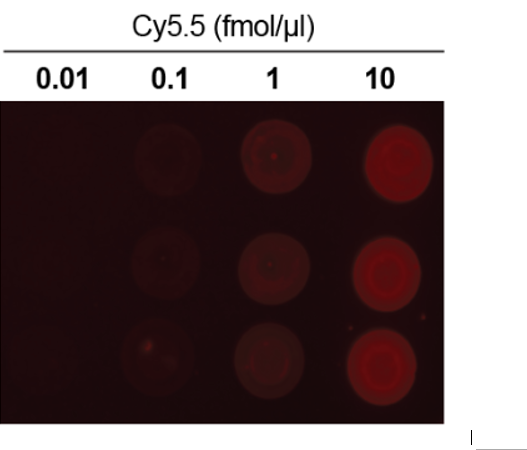

**Supplementary Figure 1.** Detection of Cy5.5 probe using Licor Odyssey. Cy5.5 probe was diluted to the mentioned concentrations and 30 µl was added onto the nitrocellulose membrane. The membrane was imaged for 10 minutes using the 700 channel.

4) Identification of a loading dye

We noticed the gel loading dye (New England Biolabs, Boston, MA, Cat#: B7024A) generated strong fluorescence on polyacrylamide gel. Thus, we examined several gel loading dyes by spotting them onto a piece of nitrocellulose membrane and imaged. Four dyes were tested: Homemade (0.25% Bromophenol blue, 0.25% xylene cyanol, 15% Ficoll type 400 in H2O), RNA dye (New England Biolabs, Boston, MA, Cat#: B0363A), Blue Juice (Invitrogen, Vilnius, Lithuania, Cat#: 10816015), gel loading dye (New England Biolabs, Boston, MA, Cat#: B7024A). As shown in Supplementary Fig. 2, the loading dye, Blue Juice generated the least amount of fluorescence. The identification of a dye that does not interfere with Cy5.5 detection is critical for the method development.

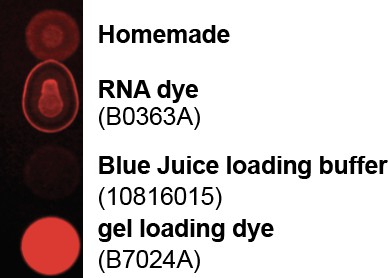

**Supplementary Figure 2.** Detection of fluorescence from dyes using Licor Odyssey. Five µl of dye was spotted onto a nitrocellulose membrane following the dilution guidelines for each dye and measured using the 700 channel by Licor Odyssey for 10 minutes.

5) Differences in fluorescence intensity of gRNA hybridization products

Fig. 1I showed large differences in gRNA signal in different RNA samples with the same amount of total cellular RNA. These differences resulted from multiple variables. The high levels of short gRNA were detected in lanes 3 and 4 because the plasmid also

expresses a Cas9 protein which is known to stabilize the gRNA. In addition, the first nucleotide of this gRNA is a G, which led to a high level of expression from the U6 promoter compared to a gRNA started with T or C^2^. The low level of long gRNA shown in lanes 5 and 6 was due to the lack of Cas9 expression from the plasmid and the first nucleotide is C. The short gRNA expression was reduced (lanes 7 and 8) because two gRNAs were expressed from the same plasmid and the first nucleotide of the short gRNA was changed to T. These results indicate that our method can measure different levels of gRNAs expression in cultured cells. This is supported by the detection of gRNA in different amounts of total RNA from cells that received viral delivery (Fig. 1J).

6) Analyzing urea effects

Urea showed great effects on the detection sensitivity because it not only suppresses the secondary structure of gRNA to allow a single hybridization to appear as a single band on the gel, but also allow the probe to more efficiently access to target gRNA. These effects are evident when total RNA hybridization with or without urea is compared, suggesting the inclusion of 4 M urea during the hybridization is important not only for suppressing the gRNA secondary structure, but also for detecting gRNA when total RNA from mammalian cells was used. To quantify the effect of urea on detection sensitivity in total cellular RNA, 20 fmol of the probe was mixed with 1 µg of total RNA from 116 cells with or without 4 M urea. Total cellular RNA was added to mimic the detection condition. We show that inclusion of urea increased the detection sensitivity by 2.5-fold (supplementary Fig. 3).

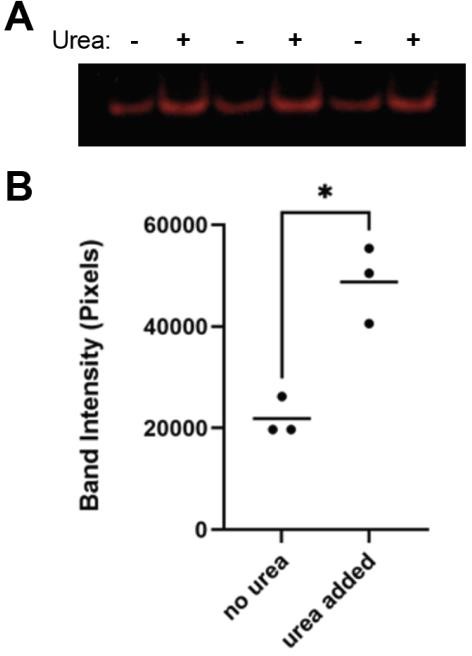

**Supplementary Figure 3.** Intensity of probe with and without urea in the presence of total RNA. **A.** 20 fmol of probe was incubated with 1 µg of total RNA with or without 4 M urea as described. **B**. The intensity of the bands was analysed using Image J (n = 3) and statistical differences were determined using an unpaired T-test. (* = p < 0.05)
